# Co-occupancy analysis reveals novel transcriptional synergies for axon growth

**DOI:** 10.1101/2020.06.12.146159

**Authors:** Ishwariya Venkatesh, Vatsal Mehra, Zimei Wang, Matthew T. Simpson, Erik Eastwood, Advaita Chakraborty, Zac Beine, Derek Gross, Michael Cabahug, Greta Olson, Murray G. Blackmore

## Abstract

Transcription factors (TFs) act as powerful levers to regulate neural physiology and can be targeted to improve cellular responses to injury or disease. Because TFs often depend on cooperative activity, a major challenge is to identify and deploy optimal sets. Here we developed a novel bioinformatics pipeline, centered on TF co-occupancy of regulatory DNA, and used it to predict factors that potentiate the effects of pro-regenerative Klf6. High content screens of neurite outgrowth identified cooperative activity by 12 candidates, and systematic testing in an animal model of corticospinal tract (CST) damage substantiated three novel instances of pairwise cooperation. Combined Klf6 and Nr5a2 drove the strongest growth, and transcriptional profiling of CST neurons identified Klf6/Nr5a2-responsive gene networks involved in macromolecule biosynthesis and DNA repair. These data identify novel TF combinations that promote enhanced CST growth, clarify the transcriptional correlates, and provide a bioinformatics roadmap to detect TF synergy.

## Introduction

As they mature, neurons in the central nervous system (CNS) decline in their capacity for robust axon growth, which broadly limits recovery from injury ^1–5^. Axon growth depends on the transcription of large networks of regeneration-associated genes (RAGs), and coaxing activation of these networks in adult CNS neurons is a major unmet goal ^2,4^. During periods of developmental axon growth, RAG expression is supported by pro-growth transcription factors (TFs) that bind to relevant promoter and enhancer regions to activate transcription ^3,6^. One factor that limits RAG expression in mature neurons is the developmental downregulation of these pro-growth TFs ^2,4^. Thus, identifying TFs that act developmentally to enable axon growth and supplying them to mature neurons is a promising approach to improve regenerative outcomes in the injured nervous system.

An ongoing challenge, however, is to decode the optimal set of factors for axon growth. To date, progress has centered on identifying individual TFs whose ectopic expression in mature neurons leads to improved axon growth. For example, we showed previously that the TF Klf6 is developmentally downregulated and that viral re-expression drives improved axon growth in mature corticospinal neurons, which are critical mediators of fine movement ^7^. The restoration of axon growth from this and other single-factor studies remains partial, however, indicating the need for additional intervention.^7–12^. One likely possibility is that because multiple TFs generally act in a coordinated manner to regulate transcription, a more complete restoration of axon growth will depend on multiple TFs. Consistent with this, regeneration competent neurons recruit groups of interacting TFs in the hours to days post-injury, which results in transcriptional remodeling leading to reactivation of growth gene networks and functional recovery ^13–15^. The need for multi-TF interventions in CNS neurons is recognized conceptually ^4,16^, but a major challenge has been to develop a systematic pipeline aimed at discovering growth-relevant TF combinations.

Here we developed a novel bioinformatics framework to detect cooperative TF promotion of axon growth. The framework centers on the concept of TF co-occupancy, in which functional interactions between TFs are detected by virtue of shared binding to common sets of regulatory DNA. We deployed the pipeline to predict novel TFs that potentially synergize with Klf6 to drive enhanced axon outgrowth. *In vitro* phenotypic screening confirmed cooperative TF activity for 12 candidates, and two independent bioinformatic analyses converged to prioritize interest in a core of three factors, Nr5a2, Rarb, and Eomes. Systematic tests of forced TF co-expression *in vivo* using a pyramidotomy model of axon injury revealed strong and consistent promotion of corticospinal tract axon growth by combined expression of Klf6 and Nr5a2. Finally, transcriptional profiling of purified corticospinal tract (CST) neurons revealed transcriptional correlates to the evoked growth, notably genes modules related to biosynthesis and DNA repair. Overall, we have identified novel combinations of transcription factors that drive enhanced axon growth in mature CNS neurons, and provide a generalized computational roadmap for the discovery of cooperative activity between TFs with potential application to a wide range of neural activities.

## Results

### In vitro screening identifies TFs that synergize with Klf6 to drive increased neurite outgrowth

Transcription factors (TFs) that functionally synergize often do so by binding DNA in close proximity and initiating complementary transcriptional mechanisms^17–20^. Thus, analysis of TF co-occupancy can reveal novel instances of cooperation between TFs^21,22^. It was shown previously that the TF Klf6 is expressed in CST neurons during developmental periods of axon growth, is down-regulated during postnatal maturation, and that forced re-expression in adult neurons modestly enhances axon growth^7^. We therefore reasoned that during development, additional TFs likely co-occupy regulatory elements alongside Klf6 and cooperate to regulate transcription. We further hypothesized that identifying these co-occupiers and supplying them to adult neurons along with Klf6 may lead to further enhancement of axon growth after injury.

To identify potential Klf6 co-occupiers, we first assembled a list of genes that decline in expression as cortical neurons mature, using data from corticospinal motor neurons (CSMNs) *in vivo* and cortical neurons ages *in vitro*^23,24^ **(Fig.1a)**. Genes common to both datasets were selected for further analysis. To tighten the focus on genes that contribute to axon growth, we selected those genes associated with Gene Ontology terms linked to axon growth, including cytoskeleton organization, microtubule-based transport, nervous system development, and regulation of growth **(Fig.1b). Supplementary Table 1** summarizes the final list of 308 developmentally downregulated pro-growth genes and their associated GO terms. For each gene, promoters were assigned as 1500 bp upstream/300 bp downstream of the transcription start site (TSS). To assign enhancers we took advantage of recent advances in machine learning-based algorithms, deploying the activity-by-contact (ABC) pipeline to reconstruct functional enhancer-gene pairs during active periods of developmental axon growth^25^. Utilizing open-access ENCODE datasets of chromatin accessibility, H3K27Ac enrichment, HiC, and expression datasets from embryonic forebrain, ABC identified a total of 1230 enhancers linked to the selected growth-relevant genes^26^ **(Fig.1c)**. Each gene was associated with between 1 and 13 enhancers (Mean 6 +/-3 SEM), 80% of which were located 10-1500 kb from the transcription start site. The full list of enhancer-gene pairs with genomic coordinates is summarized in **Supplementary Table 1**. Finally, to predict the binding of Klf6 and potential co-occupying TFs, we examined promoters and enhancers using an anchored TF motif analysis algorithm. This algorithm scans sequences for the canonical Klf6 binding motif and then identifies TF binding motifs that are over-represented in nearby DNA^27^. The result was 62 candidate TFs that were predicted to frequently co-occur with Klf6 in pro-growth promoters, pro-growth enhancers, or both **(Fig.1d)**.

We next used high content screening to test the prediction that these candidate TFs functionally synergize with Klf6 to enhance neurite outgrowth **(Fig.2a)**. Using a well-established screening platform, postnatal cortical neurons received candidate genes by plasmid electroporation and were cultured at low density on laminin substrates, followed two days later by automated tracing to quantify neurite outgrowth.^7,12,24,28–31^. Nuclear-localized EGFP served to mark transfected neurons, and βIII tubulin immunohistochemistry labeled neuronal processes for automated tracing **(Fig.2b)**. All screening plates included wells with three standard treatments: 1) EBFP plasmid alone, 2) Klf6 mixed with mCherry control and 3) Klf6 mixed with VP16-Stat3, a combination that was shown previously to elevate neurite length above Klf6 alone ^7^.This design accounts for inter-plate variability by normalizing all lengths to EBFP, while establishing on each plate both the Klf6-only effect size and the sensitivity to potential increase above that level. As expected, across the screen forced expression of Klf6 alone increased neurite outgrowth by 28% (+/-4.37% SEM) compared to EBFP control while combination with VP16-Stat3 further elevated lengths to 61% (+/-5.59%) of EBFP (p-value<.01, ANOVA with post-hoc Fisher’s LSD). The remainder of each plate was devoted to Klf6 mixed with candidate TFs, which were delivered in their native form, not modified with VP16, in order to probe endogenous activity. Each combination was tested in three separate experiments and a minimum of 150 individual cells quantified per experiment. Notably, of the 62 candidate TFs, twelve significantly elevated neurite length above the level of Klf6 alone (“hard hits”) (p-value <0.05, ANOVA with post-hoc Fisher’s LSD), and an additional five missed statistical significance but improved growth in two of three replicate experiments (“soft hits”) **(Fig.2a)**. Detailed screening data are available in **Supplementary Table 2**, and **Supplemental Figure 1** provides data for the effects of TFs on additional morphological parameters. Interestingly, all twelve TFs that elevated neurite length came from the one-third of candidates that were predicted to co-occupy with Klf6 in both promoters and enhancers, and no hits came from TFs predicted to co-occupy solely in promoters or in enhancers **(Fig.2c)**. These data support the ability of co-occupancy analysis to predict functional interactions between TFs and emphasize the importance of scanning both promoter and enhancer elements.

**Fig. 1.**
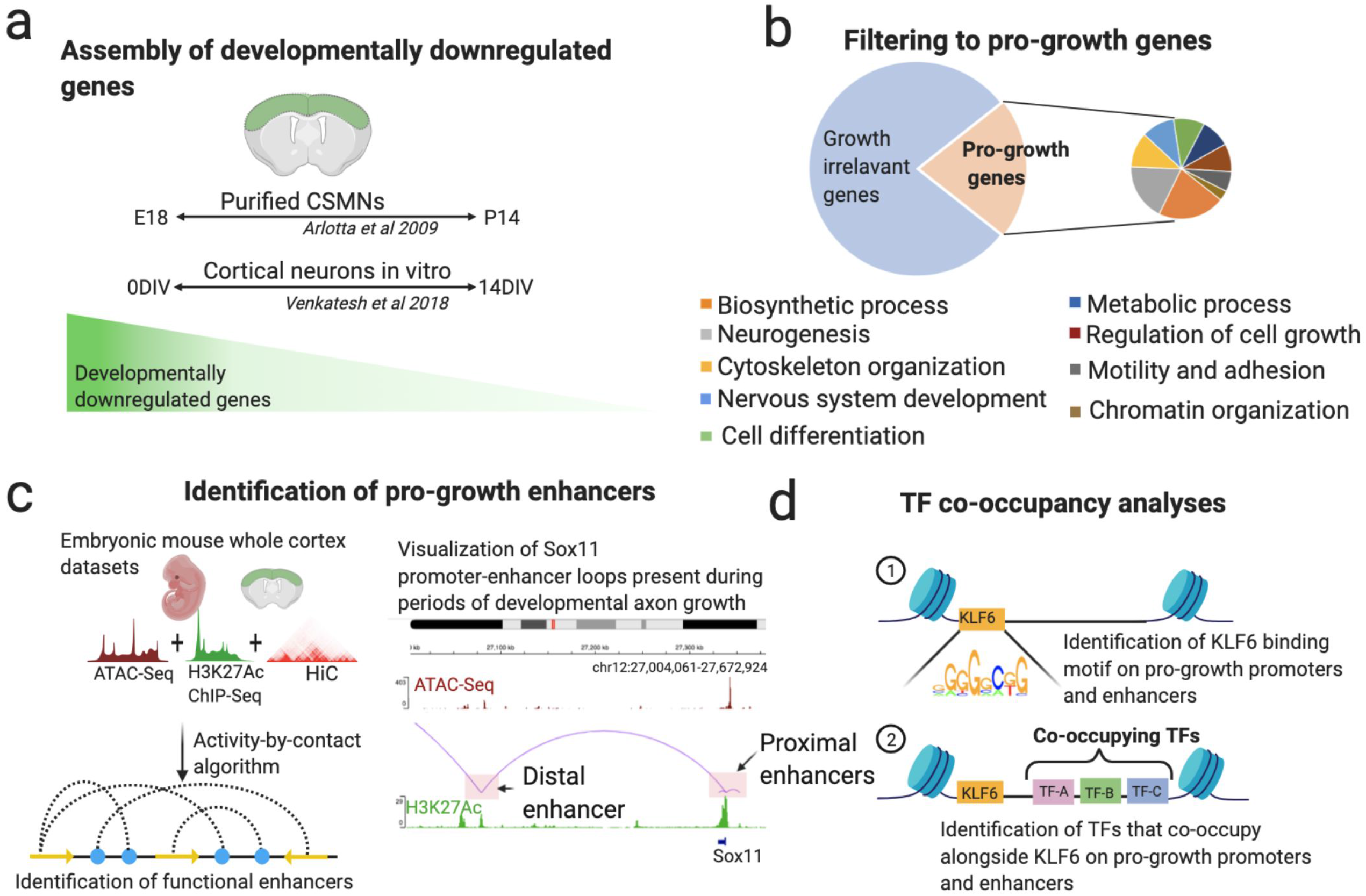
Integrated bioinformatics pipeline identifies Klf6 partner TFs likely involved in the regulation of axon growth. (a) Time-course gene expression datasets from mouse cortical neurons were integrated to isolate genes that are downregulated across cortical maturation *in vivo* and *in vitro* (b) Gene ontology analyses delineated developmentally downregulated genes that are likely growth relevant, based on functional enrichment in GO categories pertinent to axon growth (c) Activity-by-contact (ABC), an enhancer-target gene pairing algorithm, defined specific enhancer regions genome-wide that modulate the expression of growth relevant transcripts. Genome tracks are loaded to visualize the proximal and distal enhancer-promoter loops involved in the regulation of the pro-growth gene Sox11 during periods of developmental axon growth. ATAC-Seq profiles are shown in magenta and H3K27Ac profiles are shown in green. (d) TF co-occupancy analyses predicted TFs that co-occupy regulatory DNA alongside Klf6 to regulate the expression of growth-relevant transcripts.

We then used network analyses to prioritize interest in the twelve “hard hit” TFs. First, using open-access TF binding data, top transcriptional targets of the twelve hit TFs were used to generate individual TF-target gene networks. These networks were then merged, expanded to bring in immediately connected genes based on known interactions, and restructured to move genes with maximal connections to the center and minimal connections to the periphery. This resulted in a unified network with a core of TFs predicted to exert maximal influence, surrounded by shells of TFs with progressively lower connectivity (**Fig. 2c**). Notably, during the expansion step, Klf6 itself was among the genes that were spontaneously added to the network. Klf6 was the only added TF to be placed in the core, even as additional pro-growth TFs including Sox11, Stat3, Myc, and Jun appeared in inner shells. The appearance of these TFs and their central positions in the network substantiates the premise that the constructed network is Klf6-centered and relevant to axon growth^2,7–11,32–34^. Moreover, all five of the “soft hit” TFs also appeared but were placed in outer shells of the network, hinting at interactions near the threshold of screening sensitivity. Most importantly, three hit TFs (Nr5a2, Eomes, Rarb) occupied the core with Klf6, prioritizing subsequent interest. Thus, an integrated pipeline of co-occupancy analyses, high content screening, and network analyses point toward three TFs as potentially cooperative with Klf6.

**Fig. 2.**
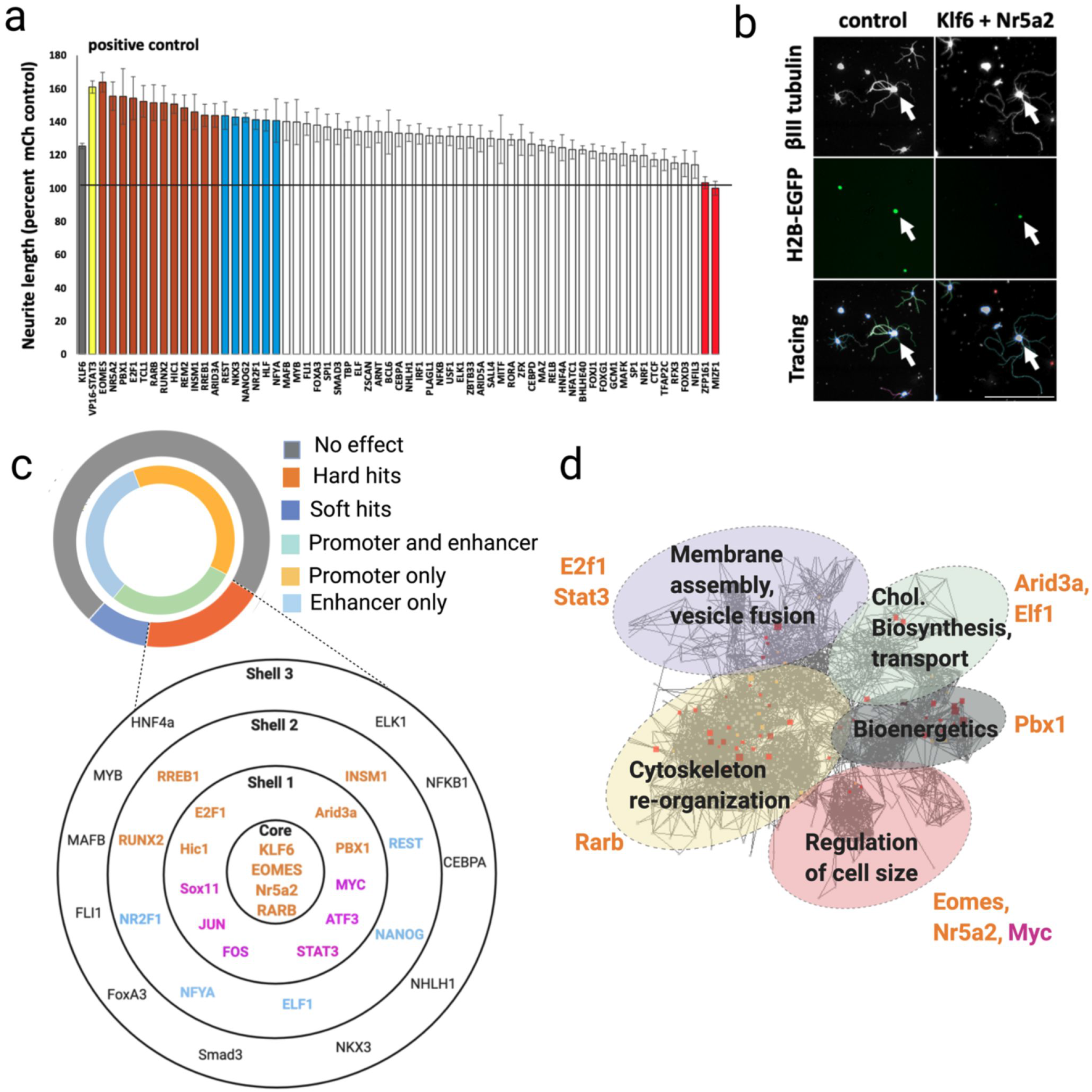
*In vitro* screening and network analyses identify a novel pro-growth transcription factor network. (a) Postnatal cortical neurons were transfected with control plasmid, Klf6 with control, Klf6 with VP16-Stat3, or Klf6 combined with 62 candidate TFs. After two days in culture, automated image analysis (Cellomics) quantified neurite lengths normalized to the average of on-plate controls. The solid black line indicates the average control length. Combined expression of Klf6 and VP16-Stat3 showed expected increase in neurite length above Klf6 alone, confirming assay sensitivity (yellow bar, p-value <0.0001, ANOVA with post-hoc Fishers). 12 candidate TFs significantly enhanced neurite outgrowth when combined with Klf6 (orange bars, p-value <0.05, ANOVA with post-hoc Fishers), and 5 more enhanced growth in 2 of 3 replicates but missed the cut-off for statistical significance (blue bars). n>150 cells in each of three replicate experiments. (b) Representative images showing transfected neurons (white arrows, H2B-EGFP) and neurite tracing. (c) Network analysis was performed on transcripts upregulated by Klf6 overexpression in culture neurons, followed by motif analysis of gene promoters within each module. Of the ten TF motifs with the highest enrichment, seven were hit TFs from the screening experiment (orange), including Rarb, Eomes, and Nr5a2. (d) Hit TFs were used to build networks. All hit TFs derived from candidates with predicted Klf6-cooccupancy in both promoters and enhancers. Rarb, Eomes, and Nr5a2 were located in the network core, where Klf6 itself also appeared during network construction. Several soft hit TFs (blue) and established pro-growth TFs (pink) spontaneously populated outer shells. Scale bar is 100 µm.

For independent validation, we performed a second analysis, this time starting from a previously published set of genes that are upregulated in cortical neurons *in vitro* upon forced expression of Klf6^7^. We again performed a network analysis of Klf6-upregulated genes, constructing functionally related subnetworks, and then examined gene promoters within each subnetwork for the enrichment of TF motifs. This independent analysis detected enrichment for the motifs of seven factors, which remarkably included the 3 “core” TFs identified above **(Fig.2d)**. Thus, genes that respond to Klf6 over-expression are enriched for the recognition motifs of Eomes, Nr5a2, and Rarb, supporting the hypothesis that they cooperate with Klf6 to activate pro-growth gene networks.

### Nr5a2 synergizes with Klf6 and Rarb to drive enhanced CST sprouting following pyramidotomy injury

We next asked whether forced expression of the three hit TFs, singly or in conjunction with Klf6, can promote the growth of CST axons *in vivo* Based on serotype availability we first verified the ability of AAV2-Retro vectors, previously shown effectively transduce CST neurons in a retrograde fashion ^35^, to also transduce neurons when applied directly to cell bodies. We delivered a titer-matched mixture of AAV-H2B-mEGFP and AAV-H2B-mScarlet to the cortex of adult mice, followed by retrograde labeling of CST neurons by cervical injection of CTB-647. Two weeks later, 3.6% (+/-1.5%) of transfected CST neurons expressed only EGFP, 1.0% (+/-0.3%) expressed only mScarlet, and 95.4% (+/-1.5%) expressed both fluorophores (**Supplementary Fig. 2)**. Accordingly, TFs were delivered to adult mice by cortical injection of Retro-AAV (hereafter shortened to AAV), with AAV-Cre acting as control and also included in single-gene treatments to equalize the viral load in all animals **(See Supplementary Table 3)**. tdTomato tracer was co-injected in serotype AAV9, avoiding potential complications of retrograde spread of the tracer itself. Mice received unilateral pyramidotomy, followed eight weeks later by quantification of cross-midline sprouting of corticospinal (CST) axons in the cervical spinal cord, measured at 200, 400, and 600□ m from the midline **(Fig.3a,d)**. Axon counts were normalized to tdTomato+ axons in the medullary pyramids (**Fig. 3b**) and injuries were confirmed by unilateral ablation of PKC□ in spinal sections (**Fig. 3c**). Raw images of all animals are provided in **Supplementary Figs. 3-5**. In addition to Cre, we also included Nkx3.2, a non-hit/non-core TF, as an additional negative control; RNAscope or immunohistochemistry confirmed overexpression of all TFs **(Supplementary Fig. 6a-e)**. As expected, Klf6 expression significantly increased normalized axon counts at all distances across the midline (p<.01, 2-way ANOVA) **(Fig.3 e,g)**. When expressed singly, none of the candidate TFs significantly elevated axon growth above the level of control **(Fig.3 e, h**,**j**,**l, n)**. When combined with Klf6, however, significant effects emerged **(Fig.3 e-o)**. Most notably, co-expression of Nr5a2 with Klf6 consistently elevated axon counts above that of Klf6 alone, and above the maximum growth in Cre controls in all animals. Co-expression of Rarb with Klf6 drove growth that exceeded Klf6 alone at 200□ m from the midline **(Fig. 3e)**. In three of nine animals, combined Eomes/Klf6 expression yielded high axon counts, but this effect was inconsistent and did not reach statistical significance **(Fig. 3e)**. Finally, as, expected, Nkx3.2 produced no significant elevation in axon growth above the level of Klf6 alone **(Fig.3 e, i)**. In summary, these data identify robust synergy between Klf6 and candidate TFs, most notably Nr5a2.

**Fig. 3.**
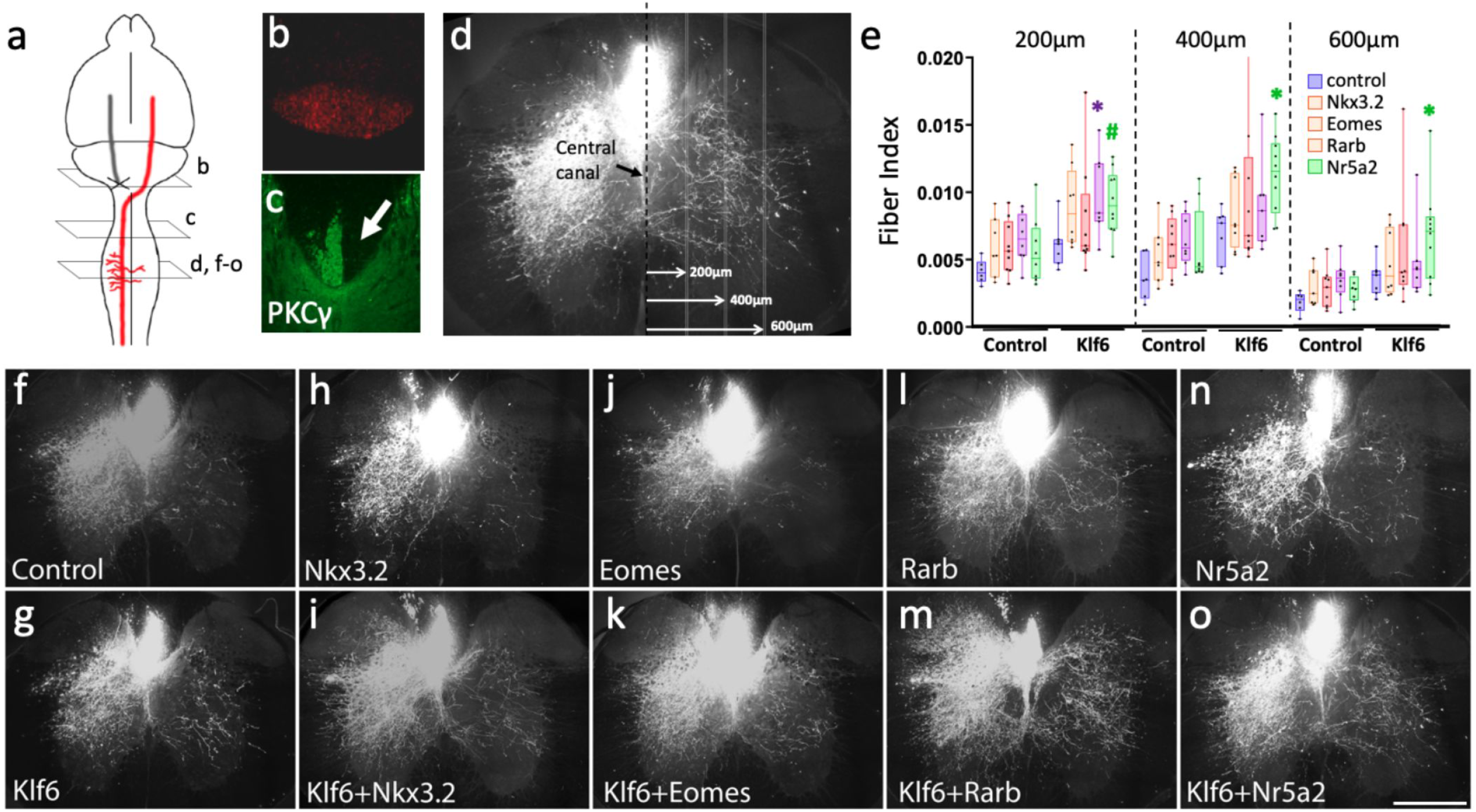
Nr5a2 synergizes with Klf6 to promote CST growth. (a) Adult mice received cortical AAV delivery of single or combinatorial TFs and contralateral pyramidotomy. (b) Co-injected tdTomato labeled CST axons, visible in cross section of the medullary pyramids. (c) PKCƳ immunohistochemistry confirmed unilateral ablation of the CST (arrow). (d,e) Cross-midline growth of CST axons was quantified in transverse sections of cervical spinal cord by counting intersections between labeled axons and virtual lines at 200µm, 400µm, 600µm from the midline, normalized to total labeled axons counted in the medulla (Fiber Index). Candidate TFs administered singly did not significantly elevate axon growth, whereas Nr5a2 and Rarb elevated growth above the level of Klf6 alone. (#p=.052, *p<.05, **p<.01, 2-way ANOVA, post-hoc Dunnett’s). (f-o) Representative images of cervical spinal cord showing elevated cross-midline growth in animals treated with Klf6 and Nr5a2. Scale bar is 500µm. n is between six and ten animals for all groups; PKCy, medulla, and spinal images from all animals are in Supplementary Figs. 2-5.

We then performed a second *in vivo* pyramidotomy experiment to confirm Nr5a2’s synergy with Klf6, and to determine whether Nr5a2 might similarly potentiate the growth of other pro-growth TFs. AAV delivery, injury, and axon quantification were identical to the first experiment, and detailed histology is available in **Supplementary Figs. 3-5**. In this experiment, Nr5a2 acted as the base gene, to which control, Klf6, or Rarb were added. Consistent with the prior experiment, combined Nr5a2/Klf6 expression elevated CST growth above the level of either alone **(Fig.4e)**. Moreover, axon growth in the lowest Klf6/Nr5a2 animal still exceeded that in the maximum control animal, illustrating the strength and consistency of the effect. Interestingly, combined Nr5a2 and Rarb also elevated CST growth above the level of either alone **(Fig.4a-e)**. This effect did not appear as striking as that of Nr5a2/Klf6, but nevertheless substantiates the ability of network analysis to predict functional TF co-operation. Thus, combinations of TFs, rationally selected based on the basis of predicted co-occupancy of pro-growth regulatory DNA, can enhance CST axon growth *in vivo*.

**Fig. 4.**
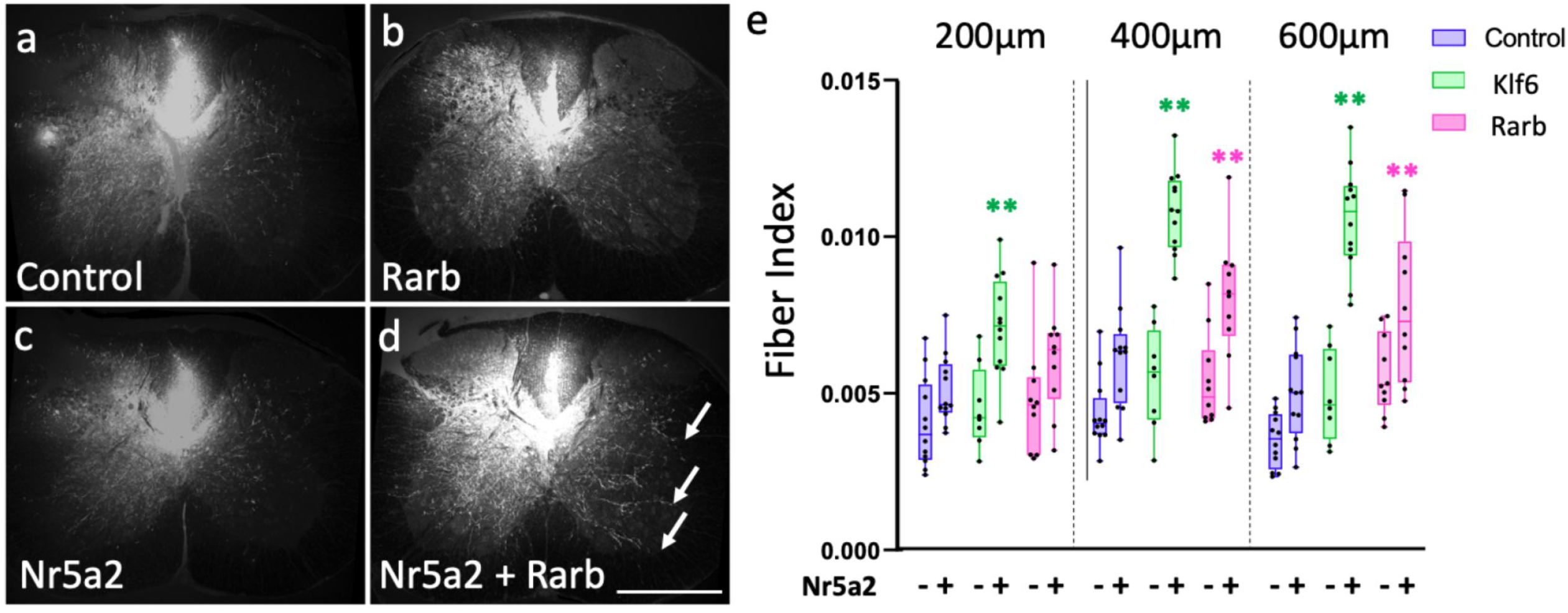
Nr5a2 synergizes with Rarb to promote CST growth. (a-d) Adult mice received cortical AAV delivery of single or combinatorial TFs and contralateral pyramidotomy. Co-injected AAV-tdTomato labeled CST axons. Elevated cross-midline sprouting of CST fiber is evident in animals treated with combined Nr5a2 and Rarb (arrows, d). (e) Cross-midline growth of CST axons was quantified in transverse spinal sections. Nr5a2 administered alone had no significant effect on axon growth (blue bars), but increased axon growth when combined with either Klf6 or Rarb, significantly above the level of either alone (green and pink bars, respectively). (**p<.01, 2-way ANOVA, post-hoc Dunnett’s). Scale bar is 500 µm. n is between 10 and 14 animals per group. PKCy, medulla, and spinal images from all animals are in Supplementary Figs. 2-5.

Next, focusing on the Klf6 and Nr5a2 combination, we performed additional experiments to substantiate and clarify the phenotypic effects *in vivo*. First, to further confirm successful co-expression we performed additional cortical injections, using the same viral loads as the pyramidotomy experiments, followed by RNAscope-based co-detection of Klf6 and Nr5a2. In animals that received AAV-tdTomato with AAV-Klf6 and AAV-Nr5a2, more than 90% of tdTomato-positive cells showed strong label with both transgenes (**Supplementary Fig. 6**). Thus tdTomato-labeled axons in the spinal cord in Nr5a2/Klf6 animals largely arose from CST neurons expressing both TFs. We next compared inflammation, gliosis, and cell death in the cortex of animals that received control versus Klf6/Nr5a2. Three days after injection, CD11b, GFAP, and TUNEL reactivity near the needle tracks were similar across treatments (**Supplementary Fig. 7a-d**,**i**,**j**). We also examined cortices in which CST neurons were identified by retrograde labeling, and again detected similarly low levels of GFAP, CD11B, and TUNEL reactivity (**Supplementary Fig. 7e-h, i, j)**. These data support a model in which Klf6/Nr5a2 act directly in CST neurons to influence axon growth, as opposed to acting indirectly by influencing gliosis, inflammation, or cell death.

We next re-examined the morphology of Klf6/Nr5a2-evoked growth by quantifying the frequency of branch points in CST axons. Starting with tissue sections from the second pyramidotomy experiment, axons that sprouted across the cervical midline were visualized at high magnification by confocal microscopy **(Fig.5a-d)**. Segments of axons that remained continuously within the plane of visualization for a minimum of 100µm were identified and traced, and the number of definitive branches along the segment was normalized to the traced length. Interestingly, CST axons in Klf6/Nr5a2-treated animals displayed significant elevation of the frequency of branch formation (**Fig. 5e**,**f, g** control: 2.5 +/-0.4 SEM branches/mm, Klf6/Nr5a2: 9.6 +/-0.5 SEM branches/mm). This finding is consistent with the *in vitro* screening data, which showed that Klf6/Nr5a2 expression increased the length of the longest neurite when branches were included in the tracing, but not when only the longest unbranched path was analyzed (**Supplementary Fig. d**,**e**). Combined, these data indicate that combined Klf6/Nr5a2 expression in cortical neurons can act to increase branch formation and growth.

**Fig. 5.**
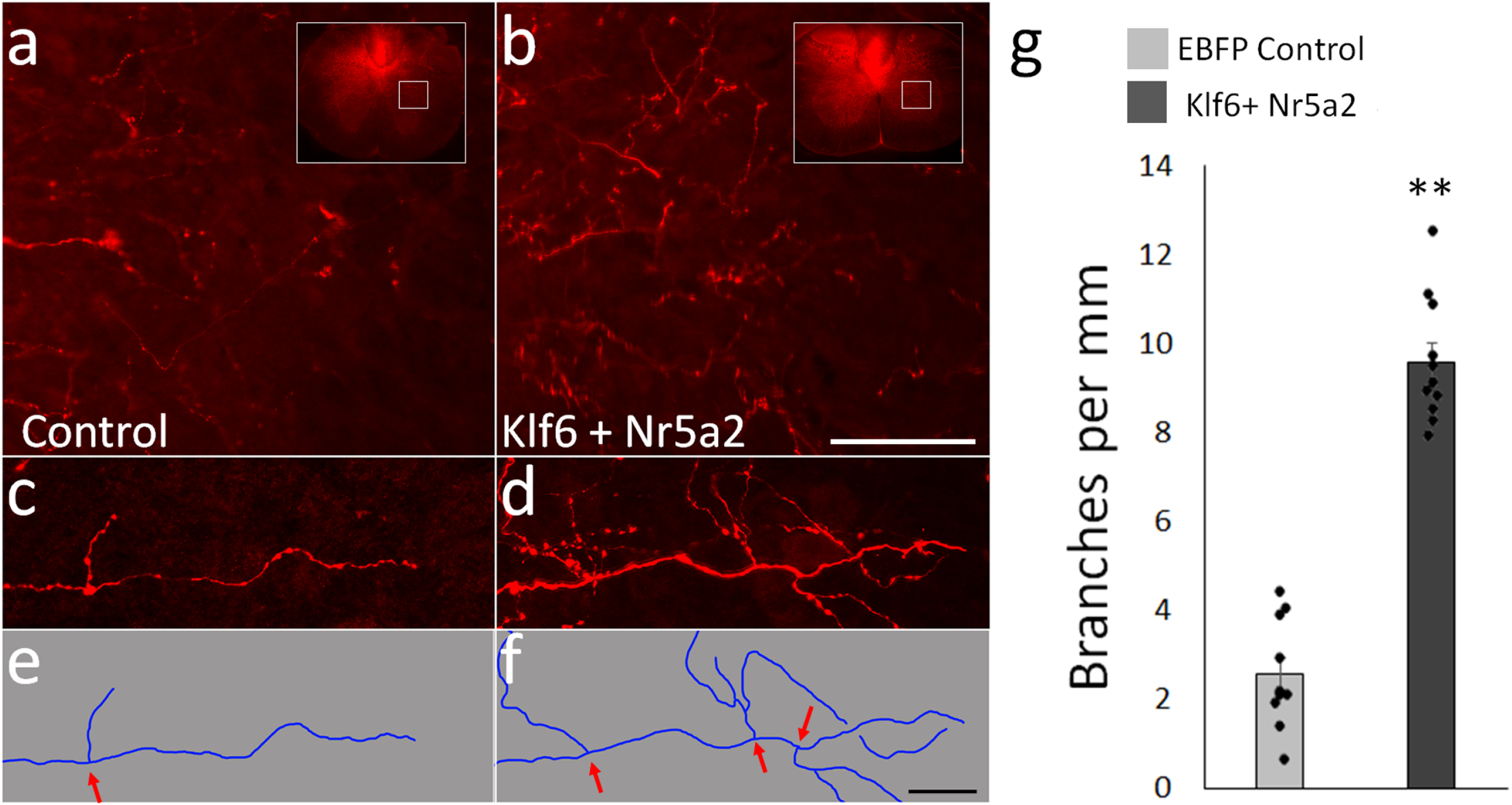
Combined Klf6/Nr5a2 treatment increases the frequency of CST axon branching. (a,b) show CST axons transduced with control (a) or Klf6 and Nr5a2 (b) that have extended across the midline into contralateral cervical spinal cord. (c-f) at higher magnification, instances of branching can be identified (arrows). (g) Animals that received combined Klf6 and Nr5a2 showed a significant increase in the frequency of branch formation above control animals (p<.001, paired t-test). A minimum of 5mm of growth, from three separate sections, were quantified for each of 10 animals in both groups. Scale bars are 100 µm (a,b) and 20 µm (c-f).

Finally, we tested the effects of forced expression of Klf6/Nr5a2 in a model of direct axon injury. Adult mice received cortical injections of AAV-EGFP tracer and either AAV-Cre control or combined AAV-Klf6 and AAV-Nr5a2. Animals received a severe crush injury to thoracic spinal cord, followed four weeks later by perfusion and visualization of axons in horizontal spinal sections (**Fig. 6a-d**). GFAP reactivity defined the injury site, which as expected spanned the entirety of the cord in all animals. In control and Klf6/Nr5a2-treated animals alike, CST axons were completely interrupted by the injury and in no animals did we observe extension of axons from the severed ends into or beyond the injury site. The distance between the lesion edge and CST axons was similar between groups, suggesting similar retraction behavior. Notably, however, axons treated with Klf6/Nr5a2 displayed a strong sprouting response into spinal tissue rostral to the injury, including robust extension across the spinal midline. Individual axons were clearly visualized crossing the midline, and Kfl6/Nr5a2 animals showed a significant elevation of axon density in contralateral spinal cord, normalized to axon counts in the medulla (**Fiber Index, Fig. 6e**). Thus, although Klf6/Nr5a2 does not confer to CST axons an ability to traverse the strongly inhibitory environment of a spinal lesion, it does act in injured axons to increase the propensity for growth by axon collaterals.

**Fig. 6.**
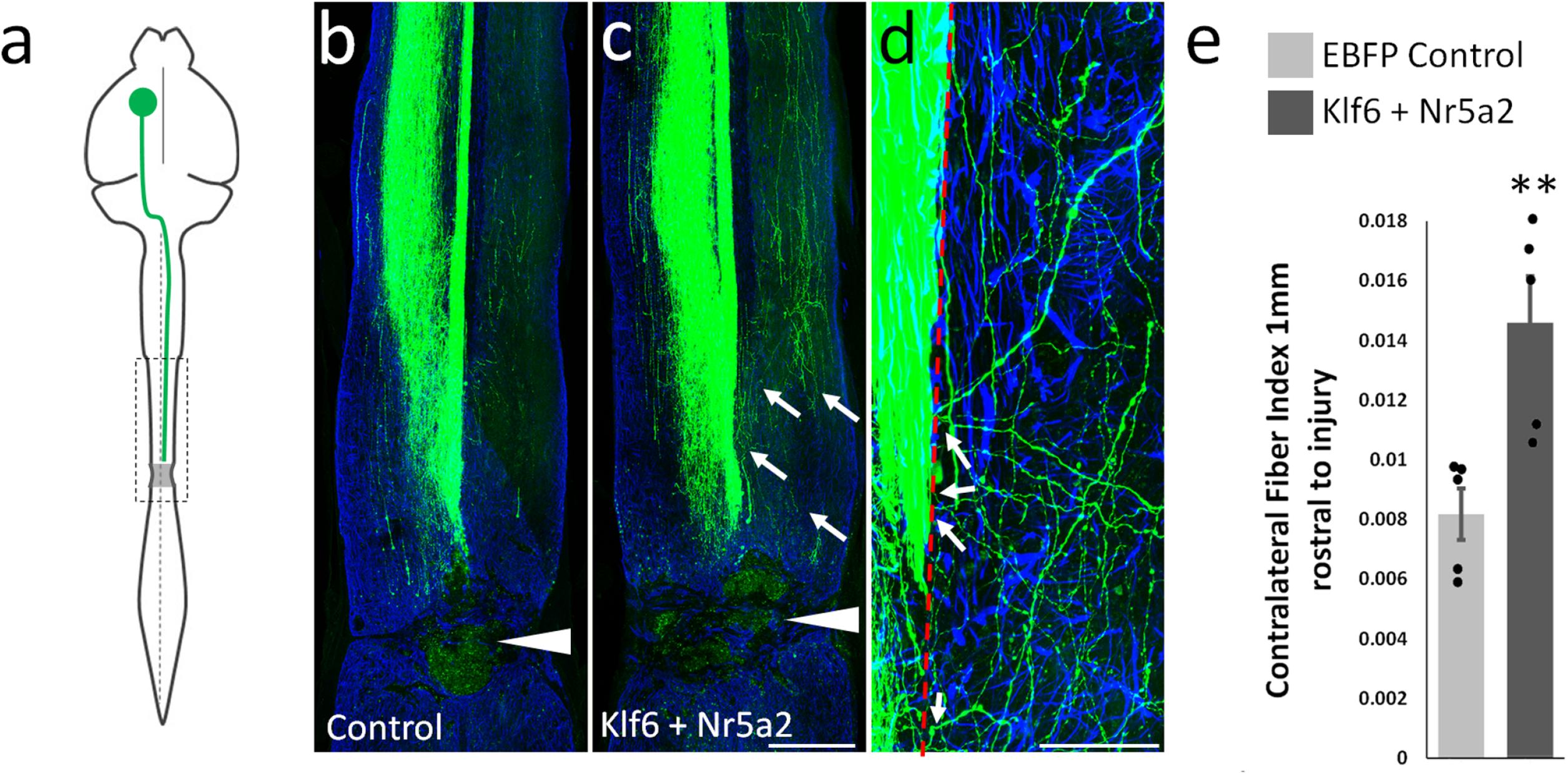
Combined expression of Klf6 and Nr5a2 in corticospinal neurons increases collateral sprouting but not extension through sites of spinal lesion. (a) Adult mice received cortical injection of AAV-EGFP tracer with AAV-Cre control or combined AAV-Klf6 and AAV-Nr5a2 and T10/11 spinal crush injuries. (b and c) show horizontal spinal sections with examples of CST axons (green), spinal injuries (arrowheads), and reactive astrocytes (GFAP, blue). Neither Cre control nor KLF6/Nr5a2-treated axons traverse the injury, but Klf6/Nr5a2-treated axons display cross-midline sprouting rostral to the injury (arrows). (d) shows CST collaterals crossing the midline (dotted line) and extending into contralateral cord. (e) Quantification of CST axons that extend 400µm from the midline, normalized to total CST axons counted in the medullary pyramids shows a significant elevation in cross-midline growth (p<.01, paired t-test, n=5 control, 5 Klf6/Nr5a2).

### Combined Klf6-Nr5a2 treatment leads to the upregulation of gene modules related to macromolecule biosynthesis and DNA repair

To probe the underlying mechanisms of growth promotion, we profiled the transcriptional consequences of single and combined expression of Klf6 and Nr5a2 in CST neurons. To isolate CST neurons, nuclear-localized fluorophores and TFs were expressed by cervical injection of Retro-AAV2, which results in highly efficient and selective transgene expression by retrograde transduction^35^ **(Fig.7a-b)**. Animals received pyramidotomy injury, followed one week later by purification of labeled nuclei of CST neurons from the spared hemisphere by flow cytometry, isolation of messenger RNA, and construction and sequencing of RNA libraries. Differential gene expression was determined using EdgeR software^36^. Compared to tdTomato control, single expression of Klf6 resulted in the upregulation of 208 transcripts (transcripts with FPKM>5 and log2fold change>1; p-value < 0.05, FDR < 0.05) **(Fig.7c)**. Reminiscent of prior findings in cultured neurons^7^, network analysis of Klf6-responsive genes revealed subnetworks with functions highly relevant to axon growth, including CNS development, neuron projection development, migration, adhesion, and cytoskeleton organization **(Fig.7d-f)**. Indeed, neuron projection and cytoskeleton networks comprised more than 60% of Klf6-responsive genes, consistent with the effects of Klf6 overexpression on axon growth **(Fig.7d-f)**. Single overexpression of Nr5a2 had little unique effects on gene expression, with only 12 transcripts identified as upregulated only in the Nr5a2 group, and 130 transcripts shared with the Klf6 treatment group **(Fig.7c)**. Dual Klf6/Nr5a2 expression, however, drove upregulation of 192 transcripts that were not significantly increased by either alone **(Supplementary Table 4, Fig.7c)**. Interestingly, network analysis of these transcripts revealed modules involved in macromolecule biosynthesis, DNA repair, and development, all of which showed substantial increases in the net expression above control levels **(Fig.7g-i)**.

**Fig. 7.**
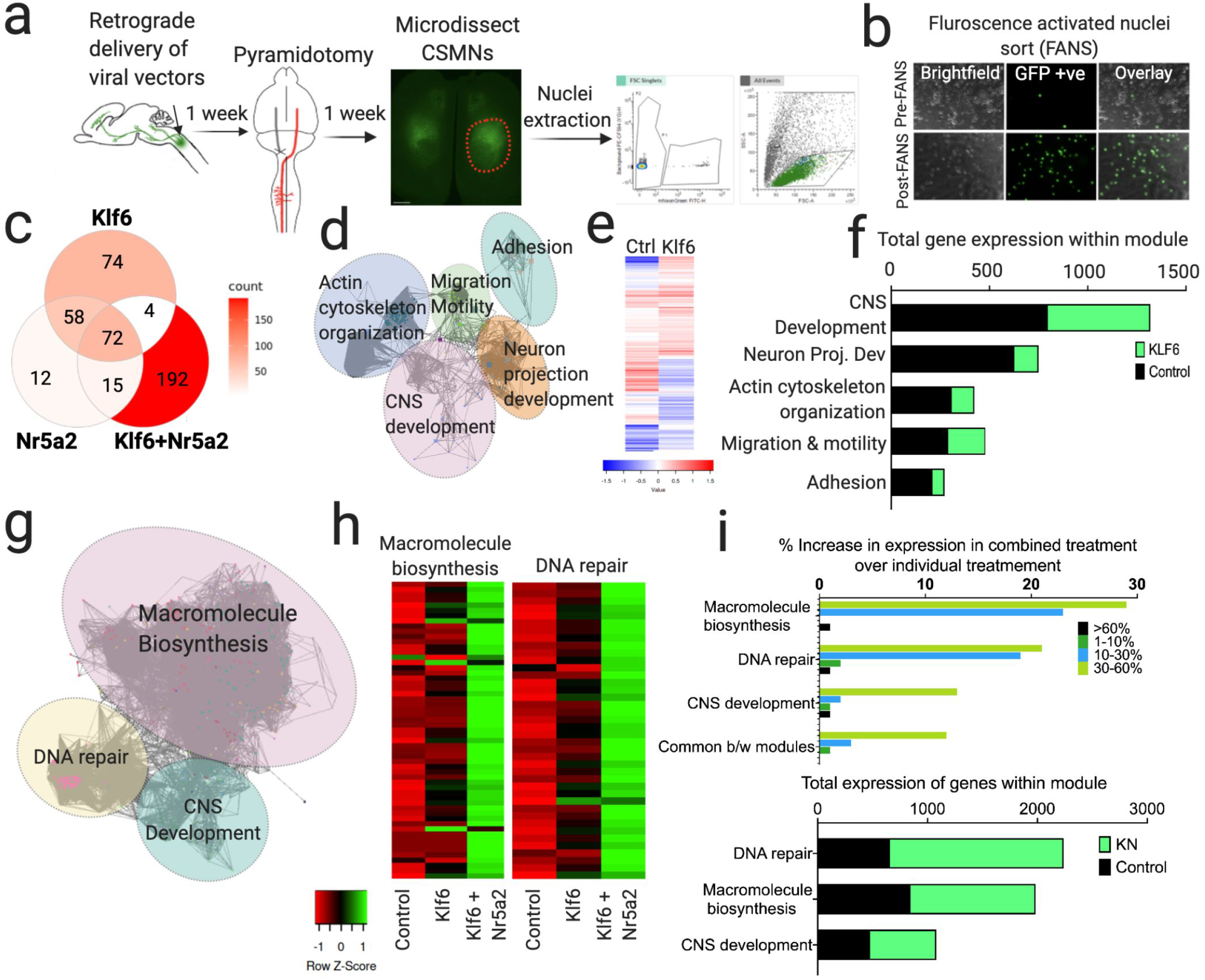
Combined Klf6, Nr5A2 treatment induces expression of gene modules involved in macromolecule biosynthesis and DNA repair. (a) Overview of sample collection for RNA-Seq analysis. (b) Shows nuclei before and after FANS, confirming successful purification of mNeongreen+ nuclei. (c) Venn diagram showing transcript comparison across the three gene treatments (d) Regulatory network analysis of genes upregulated after Klf6 overexpression revealed sub-networks enriched for distinct functional categories highly relevant to axon growth. Nodes correspond to target genes and edges to multiple modes of interaction (physical, shared upstream regulators, shared signaling pathways and inter-regulation). Only significantly enriched GO categories were included in the network analysis (p < 0.05, Right-sided hypergeometric test with Bonferroni correction, Cytoscape). (e) Heatmap showing selection of differentially expressed genes between control and Klf6 treatment. (f) shows the number of genes within each functional sub-network (top) and the total FPKM values in each in control and Klf6 overexpression groups. (g) Regulatory network analysis of genes uniquely upregulated after combined Klf6 and Nr5a2 overexpression (not upregulated in single treatments) revealed sub-networks enriched for functions in macromolecule biosynthesis and DNA repair (p < 0.05, Right-sided hypergeometric test with Bonferroni correction, Cytoscape) (h) Heatmap showing selection of differentially expressed genes in DNA repair and macromolecule biosynthesis modules. Genes that are upregulated (green) and downregulated (red) relative to control are shown. Color values indicate average FPKM expression values as shown in legend. Columns correspond to treatment groups and rows to genes. (i) indicates the percent change in fpkm values within each module, revealing that combined KLF6 and Nr5a2 treatment most affect genes in DNA repair and Macromolecule biosynthesis modules (top). Graph shows the number of genes within functional sub-networks (bottom). n=3 animals pooled per replicate, 2 reps/gene treatment.

To further validate the findings of this bulk RNA-Seq approach we performed a replicate study, this time employing a single-nuclei RNA-Seq approach. Nuclear-localized fluorophores and TFs (Control or Klf6+Nr5a2) were expressed by cervical injection of Retro-AAV2 as described above **(Supplementary Fig**.**8a)** Animals received pyramidotomy injury, followed one week later by purification of labeled nuclei of CST neurons by flow cytometry, construction and sequencing of single-nuclei libraries. Using this approach, we sequenced and analyzed ∼3000 control nuclei and Klf6/Nr5a2 treated nuclei **(Supplementary Fig**.**8b)**. The majority of nuclei partioned into clusters highly enriched for established markers of CST neurons, including Bcl11b, Cdh13, Tmem163, Crim1, Cntn6 and Slco2a1, indicating successful purification of CST neurons^23,37,38^ **(Supplementary Fig**.**8c)**. We integrated control and Kfl6/Nr5a2 single-nuclei datasets using SEURAT ^39^ and carried out differential gene expression analyses to identify Klf6/Nr5a2 responsive target genes, specifically in the high-confidence CST cluster (**Supplementary Table 4**). As expected, Klf6 and Nr5a2 themselves were detected as strongly upregulated. In addition, when comparing the output of the initial bulk RNA-Seq and single-nuclei RNA-Seq, 144 genes were commonly called as significantly regulated by Klf6/Nr5a2 in both the datasets, 101 of which were shared with the 192 transcripts called as Klf6/Nr5a2-specific in the bulk RNA-Seq experiment (>50% overla) (**Supplementary Fig 8d)**. Importantly, all shared target genes were located in clusters functionally linked to roles in Macromolecule Biosynthesis, DNA repair and CNS development, providing independent support for regulation of these processes by combined experssion of Kfl6 and Nr5a2 **(Supplementary Fig. 8e-g**). Intriguingly, recent analyses of regeneration-competent peripheral neurons also showed that genes involved in macromolecule biosynthesis and DNA repair are upregulated in the course of successful regeneration^14^, supporting their potential involvement in axon growth. In aggregate, these data substantiate the ability of forced Klf6 expression to activate gene networks relevant to axon growth and point toward the ability of dual Klf6/Nr5a2 expression to activate gene networks involved in macromolecule synthesis and DNA repair as a likely explanation for their combined enhancement of axon growth.

## Discussion

Using interlocking bioinformatics approaches, high content screening, and *in vivo* testing, we have identified new transcription factor (TF) combinations that synergize to enhance CST axon growth. In addition, transcriptional profiling and network analyses provide molecular correlates to the evoked responses. These findings clarify cellular functions that drive axon extension and serve as a roadmap for ongoing efforts to discover novel combinatorial gene treatments to promote regenerative axon growth.

Klf6 belongs to the KLF family of transcription factors, which have well-studied roles in regulating axon growth^7,12,28,40^. Retinoic acid receptor Beta (Rarb) and Nuclear Receptor Subfamily 5, Group A, Member 2 (Nr5a2/LRH-1) both belong to the nuclear receptor family of transcription factors, with diverse functions in development and cellular metabolism^41^. Rarb has previously been linked to axon growth, and a Rarb agonist is under evaluation for clinical efficacy in a brachial plexus avulsion model of injury^42^. In contrast, Nr5a2 is unstudied in the context of axon extension. It has, however, been linked to the differentiation of neural stem cells and to cellular reprogramming, and can replace Oct4 in the reprogramming of murine somatic cells to pluripotent cells^43,44^. Our present work adds to the growing body of evidence that TFs involved in neuronal differentiation may positively influence axon growth^9,13^.

An important finding is that although Nr5a2 does not improve axon growth when expressed individually, it strongly potentiates the effect of overexpressed Klf6. The importance of combinatorial gene activity for axon growth, as opposed to single factor effects, is increasingly recognized ^2,4,45^. Regeneration-competent cell types such as zebrafish retinal ganglion cells and peripherally injured sensory neurons respond to axotomy by upregulating groups of TFs that likely synergize to drive regenerative axon growth ^14,15,46^. An unmet challenge, however, has been to turn the conceptual appreciation for the role of TF synergy into an operational workflow to discover effective gene combinations for axon extension in the injured nervous system. Whereas prior efforts have focused on direct protein-protein interactions to predict TF synergy^46^, here we exploited the concept of TF co-occupancy. A search for factors with recognition motifs in proximity to those of Klf6 yielded a set of 62 candidate TFs, which was further narrowed by phenotypic screening. A key observation from the cell culture experiments is that none of the hit TFs were predicted to co-occupy solely in promoter regions; rather, all hit TFs were characterized by predicted co-occupancy in both promoters and enhancers. This suggests a key role for enhancer elements in regulating pro-growth gene transcription, in line with the recent finding in DRG neurons that modulation of enhancers may be critical for transcriptional activation of growth genes^13,47^.

Finally, guided by network analysis of the screening results, a systematic campaign of combinatorial testing *in vivo* identified highly consistent enhancement of CST axon growth by combined expression of Klf6 and Nr5a2. Importantly, in a model of pyramidotomy injury delivery of Kfl6 and Nr5a2 did not affect cell survival, but did affect both the overall density and frequency of branching of collateral axons in contralateral spinal cord. Moreover, in axotomized CST axons, forced expression also increased the growth of collateral branches, although not regenerative growth across a spinal lesion. Importantly, in contrast to prior reports of CST regeneration through crush sites, injuries here were performed using double crushes with wider forceps, thus producing complete lesions without astrocytic bridges^48,49^. The absence of regenerative advance in this model indicates that treated axons remained sensitive to environmental inhibition at the lesion, but do not rule out the possibility that Klf6 and Nr5a2 treatment could produce regenerative growth into more favorable environments, for example progenitor-derived tissue grafts^50^. In addition, the strong effects of combined Kfl6 and Nr5a2 on collateral growth in both spared and injured CST axons suggests that this treatment could be highly beneficial in situations of partial injury by enhancing increased branch elaboration by spared axons and potentially fostering relay circuity by injured axons^51,52^. A critical question for future research is whether the stimulated branches succeed in forming functional synapses on target cells in the spinal cord. To our knowledge, this is the first evidence of TF synergy in driving growth of axons following CNS injury *in vivo*. Although these experiments were anchored on Klf6, these data show a pipeline that can be re-deployed to identify additional TF combinations that synergize to drive growth, or potentially other cellular functions. To our knowledge, this is the first evidence of TF synergy in driving growth of axons following CNS injury *in vivo*. Although these experiments were anchored on Klf6, these data show a pipeline that can be re-deployed to identify additional TF combinations that synergize to drive growth, or potentially other cellular functions.

A key insight from the current work is to clarify *in vivo*, in purified CST neurons, the transcriptional events that are triggered by Klf6, Nr5a2, and the two together. Forced expression of Klf6 activates highly interconnected groups of genes with functions in various aspects of axon growth spanning terms such as Neuron Projection Development, Cytoskeleton organization, Motility, and Adhesion. We have previously shown that gene modules involved in Cytoskeleton remodeling and motility are evoked in response to Klf6 overexpression in cortical neurons *in vitro*^7^, consistent with our *in vivo* findings here. This overlap underscores the utility of *in vitro* screening approaches in delineating transcriptional networks relevant to axon growth. Importantly, this finding illustrates the potential of TF-based interventions to trigger broad changes in gene expression; it is most likely that improvements in axon growth reflect the net change in a wide set of transcripts, as opposed to acting through any single target. Combined Klf6/Nr5a2 expression, which led to large and consistent increases in axon growth above those triggered by Klf6 alone, also caused the upregulation of unique modules of genes with functions in macromolecule biosynthesis and DNA repair. These modules were activated by neither Klf6 nor Nr5a2 alone, suggesting that enhanced biosynthesis and/or DNA repair contribute to the enhanced growth in Klf6/Nr5a2-treated animals. A role for the biosynthesis of macromolecules is highly plausible, as active growth depends on new cellular material. Indeed, regeneration-competent zebrafish RGCs and mammalian DRGs also respond to injury by upregulating genes involved in macromolecule biosynthesis^15,13,14^. Moreover, a very recent study in DRG neurons found that peripheral injury, which triggers axon growth, resulted in a sustained upregulation of genes involved in biosynthesis. In contrast, a central injury that does not trigger regeneration led to eventual downregulation, suggesting that a reduction in biosynthesis pathways may restrict axon growth after spinal injury^53^. Our new findings regarding the effects of Klf6/Nr5a2 add to multiple lines of evidence that the upregulation of genes involved in biosynthesis may be an evolutionarily conserved molecular signature of neurons mounting a successful growth response.

Regarding DNA repair pathways, it is interesting to note that increased cellular metabolism and transcription frequently lead to DNA damage^54,55^. Moreover, disruption of DNA repair machinery in mouse retinal progenitors or cortical progenitors leads to reduced, disturbed trajectories of axon growth and guidance^56,57^. Thus during development, proper axon growth may require efficient DNA repair machinery. Similarly, peripherally injured DRG neurons upregulate several DNA damage response marker genes, and blocking this response leads to reduced neurite outgrowth *in vitro* and impaired regeneration *in vivo*^58^. The finding that combined Klf6/Nr5a2 expression activates genes involved in DNA repair suggests the intriguing hypothesis that this repair machinery may aid successful axon regrowth in CNS neurons; future work can address this notion.

In summary, a new bioinformatic and screening platform, centered on the concept of TF co-occupancy, has revealed synergy between transcription factors Klf6 and Nr5a2 that leads to improved axon growth after CNS injury.

## Methods

### Identification of developmentally downregulated, growth-relevant transcripts

All expression datasets used to identify developmentally down-regulated genes are deposited at NCBI GEO (In vivo CSMNs data by^23^ - GSE2039; In vitro primary neurons aged in culture by^24^, SRP151916). *In vitro* RNA-Seq datasets were processed as described previously ^24^. Trimmed reads were mapped to the rat reference genome [UCSC, Rat genome assembly: Rnor_6.0] using HISAT2 aligner software (unspliced mode along with–qc filter argument to remove low quality reads prior to transcript assembly)^59^. Transcript assembly was performed using Stringtie ^60^ and assessed through visualization on UCSC Genome Browser. Differential gene expression analysis was performed using Ballgown software (Default parameters) ^60^. Transcripts were considered significant if they met the statistical requirement of having a corrected p-value of <0.05, FDR <0.05.

For the *in vivo* CSMN microarray datasets, weighted correlation network analysis (WGCNA) was performed as described previously ^61^. First, relevant microarray datasets (GSE2039) were loaded into R and basic pre-processing of data was performed to remove outlier data and handle probes with missed data. Outlier data was identified by performing hierarchical clustering to detect array outliers. Numbers of missing samples in each probe profile were counted and probes with extensive numbers of missing samples were removed. All pre-processing steps were performed as described in ^61^. Next, a weighted correlation network was constructed by creating a pairwise pearson correlation matrix, which was transformed into an adjacency matrix using a power of 10. Then, topological overlap was calculated to measure network interconnectedness and average linkage hierarchical clustering was done to group genes on the basis of the topological overlap dissimilarity measure. Finally, using a dynamic tree-cutting algorithm we identified 20 gene modules. Gene modules with developmental down-regelation in expression of 2-fold and above were isolated for ontology analyses. Gene ontology analyses were performed using Database for Annotation, Visualization and Integrated Discovery (DAVID) Bioinformatics Resource^62^ and genes with terms relevant to axon growth (FDR<0.05) based on literature review were user-selected for further analyses.

### Enhancer Identification

Enhancers relevant to pro-growth genes were identified by running the Activity-by-contact (ABC) algorithm^25^ using code described here - https://github.com/broadinstitute/ABC-Enhancer-Gene-Prediction. ENCODE datasets used for the analysis are - ATAC-Seq (ENCSR310MLB, ENCSR836PUC), H3K27Ac histone ChIP-seq (ENCSR094TTT, ENCSR428OEK), RNA-seq (ENCSR362AIZ, ENCSR080EVZ) and HiC (GSE96107). The algorithm has three sequential steps – definition of candidate enhancers, quantification of enhancer activity and calculation of ABC scores. For definition of candidate enhancers, indexed and sorted bam files of ATAC-Seq and H3K27Ac ChIP-seq datasets were supplied and peaks were called using MACS2, with the argument --nStrongestPeaks 15000. For quantification of enhancer activity, reads counts were calculated using peak files processed in the previous step to yield a final list of candidate enhancer regions with ATAC-seq and H3K27ac ChIP-seq read counts within gene bodies and promoters. Finally, ABC scores were calculated using arguments --hic_resolution 5000, - scale_hic_using_powerlaw and a threshold of 0.01. The default threshold of 0.01 corresponds to approximately 70% recall and 60% precision. All enhancer-gene pairs that passed the above threshold were used for subsequent analyses.

### Co-occupancy motif analyses

Transcription factor binding site/motif analysis on pro-growth gene promoters and enhancers was performed using opossum v3.0 software^27^. For promoter analyses, the list of pro-growth genes was supplied along with promoter co-ordinates to be used for scanning (upstream/downstream of TSS – 1000/300 bps). For enhancer motif analyses, a unified file containing a list of FASTA formatted sequences corresponding to pro-growth enhancer regions was supplied.

Mouse anchored TF-cluster analyses were performed using search parameters – JASPAR CORE profiles that scan all vertebrate profiles with a minimum specificity of 8 bits, conservation cut-off of 0.40, matrix score threshold of 90%, and results sorted by Z-score >=10. TFs were sorted into three categories – TFs with over-represented sites within pro-growth promoters, TFs with over-represented sites within pro-growth enhancers and TFs with over-represented binding sites within both pro-growth promoters and enhancers.

### Network analyses

Hit TF-target gene networks were constructed using ENCODE ChIP-Seq data for hit TFs using the Harmonizome database^63^. For each of the 12 hit TFs, TF target gene lists were retrieved from Harmonizome individually and merged before visualization on the Cytoscape platform (v3.7.1). Cytoscape plug-ins ClueGO and CluePedia were used for network visualization. iRegulon plug-in on the Cytoscape platform was used to expand the network one-level and bring in additional regulators. Next, we calculated connectivity scores to assign centrality using the Network Analyzer option within Cytoscape 3.7.1. TFs were ranked in descending order of connectivity scores to assign centrality. Core TFs were those TFs that had the highest connectivity scores and were centrally located in the network, followed by TFs with decreasing connectivity scores that occupied peripheral positions in the network. Upstream regulator analysis on Klf6 target genes was run on differentially expressed genes listed in^7^ using motif analyses described above. Motif analyses were done in batches such that each batch had genes belonging to one functional sub-network to identify Klf6 synergizers by functional category.

### Cloning and virus production

Constructs for candidate genes were purchased from Dharmacon or Origene, and relevant accession numbers for all 62 candidate TFs are summarized in **Supplementary Table 2**. mScarlet was a gift from Dorus Gadella (RRID:Addgene_85044), pAAV-CAG-tdTomato (codon diversified) was a gift from Edward Boyden (RRID:Addgene_59462), and mNeonGreen sequences were synthesized by Genscript, based on the amino acid sequence of ^64^. For viral production, genes were cloned into an AAV-CAG backbone (Addgene-Plasmid #59462) using standard PCR amplification, as described in ^12^. Maxipreps were prepared by Qiagen Endo-free kits and fully sequenced, and AAV9-tdTomato (Addgene Plasmid #59462) and AAV2-Retro of all other constructs were produced at the University of North Carolina Viral Vector Core and brought to 1×10^13^ particles per ml in sterile saline prior to injection. Viruses were mixed at a 3:3:2 ratio of combinatorial test genes and tracer; viral treatments are detailed in **Supplementary Table 3**.

### Cortical cell culture and analysis of neurite outgrowth

All animal procedures were approved by the Marquette University Institutional Animal Care and Use Committee. Cortical neurons were prepared from early postnatal (P5-P7) Sprague Dawley rat pups (Harlan), with each experimental replicate derived from a separate litter. Procedures for dissociation, transfection, immunohistochemistry, imaging, and neurite outgrowth analysis were performed as in^7^. Plasmid expressing nuclear localized EGFP, mixed with text plasmids at a 1:3 ratio, served to identify transfected cells. Each 24-well culture plate included three standard treatments: EBFP control, Klf6 plasmid mixed in equal parts with mCherry control, and Klf6 mixed in equal parts with VP16-Stat3. Remaining wells contained Klf6 mixed with candidate TFs. All morphological values were normalized to within-plate EBFP control values, the Klf6-only treatment served to set the level of Klf6’s individual effect, and the Klf6/VP16-stat3 combination confirmed sensitivity of the assay plate to TF synergy ^7^. For all cell culture experiments, neurite length from a minimum of 150 cells per treatment was averaged, and each experiment was repeated a minimum of three times on separate days. These averaged values were the basis for ANOVA with Fisher’s multiple comparisons. Neurite outgrowth values, number of cells analyzed for every screening experiment are summarized in **Supplementary Table 2**.

### Viral delivery to cortical neurons and pyramidotomy injuries

*In vivo* experiments were performed in a double-blind fashion, with non-involved lab personnel maintaining blinding keys. Experiments used adult C57BL/6 mice, 8 to 10 weeks of age at the start of the experiment. The first pyramidotomy experiment used female mice, and the second pyramidotomy and spinal crush experiment used mixed sex at approximately equal numbers; sex had no significant effect on axon growth in either experiment (p>.05, 2-Way ANOVA). Animals were randomized prior to viral treatment, with each surgical day including equal numbers from each group. Cortical neurons were transduced using intracerebral microinjection as described in^7,8^. Briefly, mice were anesthetized with ketamine/xylazine (100/10 mg/kg, IP), mounted in a stereotactic frame, and skull exposed and scraped away with a scalpel blade. 0.5 μl of virus particles were delivered at two sites located 0 mm / 1.3 mm and 0.5 mm / 1.3mm (anterior / lateral from Bregma) in pyramidotomy experiments, and -0.6 mm / 1.3 mm and -1.2 mm / 1.3mm (anterior/lateral from Bregma) in spinal crush experiments, at a depth of 0.55 mm, and at a rate of 0.05 ul/min using a pulled glass micropipette connected to a 10 µl Hamilton syringe driven by a programmable pump (Stoelting QSI), with one minute dwell time. For spinal injections, mice were mounted in a custom spine stabilizer and viral particles or Cholera Toxin Subunit B conjugated to Alexa Fluor 647 (CTB-647) in sterile 0.9% NaCl (C22841-Thermofisher, Waltham, MA, final concentration of 2%) were injected to the spinal cord through a pulled glass micropipette fitted to a 10 μl Hamilton syringe driven by a Stoelting QSI pump (catalog # 53311) (pumping rate:0.04 µl/min) between C4 and C5 vertebrae, 0.35 mm lateral to the midline, and to depths of 0.6 mm and 0.8 mm, 0.5 µl at each depth. Unilateral pyramidotomy was performed as described in^8,12^. Briefly, a ventral midline incision was made to expose the occipital bone, the ventrocaudal part of which was removed using fine rongeurs. The dura was punctured, and the right pyramid cut completely using a micro feather scalpel. For spinal crush injuries, mice were anesthetized and mounted in a spine stabilization device, a laminectomy performed at T12 vertebrae / T10/11 spinal cord, and forceps of width 250 µm used to crush the cord for ten seconds, then grip-reversed and repeated for another ten seconds.

### RNAScope

RNAscope kits are commercially available from Advanced Cell Diagnostics (ACD). We utilized the Multiplex v2 system for all RNAScope experiments (Cat-323100,ACD). Mouse brains were snap-frozen on dry ice and cryosectioned (25µm). Slides were baked at 60°C for 40 mins followed by RNAScope procedures according to manufacturer’s instructions. Briefly, hydrogen peroxide solution was applied to baked slices and incubated for 10 mins at RT. Next, slices were incubated in boiling antigen retrieval solution (<98 °C) for 5 mins. Following retrieval, slices were washed in Nuclease free water three times, 5 mins per wash. Then, brain slices were dehydrated in 100% ethanol briefly followed by treatment with ProteaseIII for 20 mins at 40°C. Following protease incubation, slices were washed in Nuclease free water three times, 5 mins per wash and probes were applied and allowed to incubate for 2 hours at 40°C. The following probes were purchased off the catalog-RNAscope® Probe - Mm-Klf6 (Cat-426901, ACD), RNAscope® Probe - Mm-Nkx3-2 (Cat-526401, ACD),RNAscope® Probe - Mm-Rarb (Cat-463101, ACD). We designed a custom probe for Nr5a2, targeting a region common to all variants(RNAscope® Probe - Mm-Nr5a2-O1-C2, Cat-547841-C2, ACD). All all probes were detected with TSA plus fluorophore used at 1:750 dilution. Before mounting the slices, a brief 5 minute incubation with DAPI was performed to label the nuclei. All RNAScope experiments were carried out between 1-3 days post cryosectioning to ensure tissue integrity.

### Immunohistochemistry

Adult animals were perfused with 4% paraformaldehyde (PFA) in 1X-PBS (15710-Electron Microscopy Sciences, Hatfield, PA), brains, and spinal cords removed, and post-fixed overnight in 4% PFA. Transverse sections of the spinal cord or cortex were embedded in 12% gelatin in 1X-PBS (G2500-Sigma Aldrich, St.Louis, MO) and cut via Vibratome to yield 100 µm sections. Sections were incubated overnight with primary antibodies PKCγ (SC C-19, Santa Cruz, Dallas, TX, 1:500, RRID: AB_632234), GFAP (DAKO, Z0334 1:500, RRID:AB_10013482), or Cd11b (Invitrogen 14-01120-82 1:500, RRID:AB_2536484) rinsed and then incubated for two hours with appropriate Alexa Fluor-conjugated secondary antibodies (Thermofisher, Waltham, MA, 1:500.) For TUNEL staining (In Situ Cell Death Kit, Roche), fresh frozen transverse cryostat sections (30 μm) were post-fixed in 4% PFA 15 minutes, incubated in ethanol/acetic acid for 10 minutes, 0.4% Triton for 30 minutes, and with probe mixture for 1 hour^8^. Staurosporine control (1 μl, 1 mM; Sigma) was cortically injected 2 days prior to sacrifice. Fluorescent images were acquired using Olympus IX81 or Zeiss 880LSM microscopes.

### Quantification of axon growth

In pyramidotomy experiments, axon growth was quantified from four 100 µm transverse sections of the spinal cord of each animal that spanned C2 to C6 spinal cord. Starting from a complete series of transverse sections, the rostral-most was selected from C2 as identified by the morphology of the ventral horns of the spinal cord, and the next three selected at 1.6mm intervals in the caudal direction. Each section was imaged on Nikon confocal microscope (Nikon AR1+) using a 20X Plan Apochromatic (MRD00205, NA 0,75) objective, gathering 20 µm of images at 5 µm intervals and then creating a maximum intensity projection. On the resulting image, virtual lines of 10 µm width were placed at 200, 400, and 600 µm from the midline and intersection of tdTomato+ axons with these lines were quantified. For normalization, 100 µm transverse sections of the medullary pyramids were prepared by vibratome sectioning and imaged on a Nikon confocal microscope (Nikon AR1+) using a 60X Apochromatic Oil DIC N2 (MRD71600, NA 1.4) objective. Seven virtual lines of 10 µm width were distributed evenly across the medial/lateral axis of the medullary pyramid, the number of tdTomato+ axons visible in each was determined, and the total number of axons extrapolated based on the area of the entire pyramid and the area of the sampled regions. Fiber index was calculated as the average number of axons at each distance from the midline across the 4 replicate spinal sections, divided by the calculated number of axons detected in the pyramid. Counting of digital images was performed by three blinded observers, with final values reflecting the average. Exclusion criteria for pyramidotomy experiments were animals with less than 80% decrease in PKCγ in the affected CST. For spinal crush injuries, quantification and normalization of cross-midline sprouting was nearly identical to those for pyramidotomy, with the exception that quantification was performed in 4 horizontal sections of spinal cord, with the sampling area set 400 µm from the midline. To quantify axon branching, the following procedure was employed. First, in transverse sections of cervical spinal cord, tdTomato+ CST axons were identified in a sampling region between 200 and 600 µm from the midline. Next, axon segments traced, with the requirement that sampled segments must remain in a single confocal imaging plane at 60X magnification for a minimum of 100 µm traced length. Finally, using Z-stacks to definitively distinguish true branches from near-plane intersections, the number of branches was counted in each sampled segment. A minimum of 5mm of total length, from three separate spinal sections, were sampled for each animal.

### Fluorescence activated nuclei sorting (FANS)

Equal numbers of male and female mice were used in every experiment in concordance with NIH guidelines. Adult mice received retrograde injections of viral vectors for Ctrl treatment, Klf6 alone treatment, or combined Klf6+ Nr5a2 treatment. One week later, animals were challenged with pyramidotomy injuries, as described above. One week post-injury, animals were euthanized, and the motor cortices were dissected. Tissue was collected only from the hemisphere contralateral to the pyramidotomy injury, thus isolating spared neurons and not neurons that experienced direct axotomy. Dissected cortices were minced finely using razor blades and transferred to pre-chilled 15 ml Dounce homogenizer filled with 3 ml Nuclear release buffer (320mM Sucrose, 5mM Cacl2, 3mM MgCl2, 10mM Tris-HCL, 0.3% Igepal). Tissue was dounced 15X while on ice and filtered sequentially via a 50um, 20um filter, and used as input for flow cytometry. Dissociated nuclei were flow-sorted on a BD FACS Melody using an 80um nozzle and a sequential gating strategy (Sort type: Purity). Specifically, nuclei were first gated by SSH-H vs FSC-H and SSC-A vs FSC-A to separate debris vs intact nuclei. Nuclei that passed these filters were then filtered by SSC-W vs SSC-H to eliminate potential doublets. Finally, nuclei were gated by levels of fluorescence marker such that only the brightest nuclei are collected to a goal of approximately 30,000 events for RNA-Seq and 4000 events for single-nuclei RNA-Seq.

### RNA-Seq data generation and analysis

Following FANS, nuclei were sorted directly into Trizol, lysed, and RNA was extracted according to manufacturer’s instructions. Only samples with a RIN score of 8 and above were used for library prep. Total RNA was used for library prep using Takara SMARTer stranded total RNA-Seq kit v2 according to the manufacturer’s instructions. Samples were sequenced at the University of Wisconsin-Madison Genomics core to a depth of 25 million paired-end reads on an Illumina HiSeq platform, with two replicates per treatment. Raw FASTQ files were fed into ENCODE consortia RNA-Seq pipeline described here - https://github.com/ENCODE-DCC/rna-seq-pipeline. Read alignment was performed using STAR^65^ and transcript quantification performed using Kallisto^66^. Transcript counts from Kallisto were used for performing differential gene expression (DEG) analyses using EdgeR^36^. Transcripts were considered significant if they met the statistical requirement of having a corrected p-value of <0.05, FDR <0.05. Differentially expressed genes were further filtered to only include transcripts with FPKM > 5 and log2Foldchange > 1. RNA-Seq datasets have been deposited with NCBI GEO (SRP151916). Network analysis on significantly differentially expressed genes (upregulated) was performed on the cytoscape platform v3.7.1 using plug-ins clueGo and CluePedia. Functional analysis mode was selected and the following GO categories were selected for analyses-GO Biological Process, KEGG pathways, Reactome Pathways and Wiki pathways. Network specificity was set to medium and GO tree interval was set between 3-8. GO term kappa score was set to default values (0.4) and leading group term was set to rank by highest significance. Lastly, GO term fusion was also selected and only sub-networks with significantly enriched GO terms were used to generate network visualizations (p < 0.05, Right-sided hypergeometric test with Bonferroni correction, Cytoscape).Expression heat-maps were assembled using heatmapper.ca using average linkage and Euclidean distance parameters for clustering.

### Single nuclei RNA-Seq data generation and analysis

Equal numbers of male and female mice were used in every experiment in concordance with NIH guidelines. Adult mice (8 weeks) received retrograde injections of viral vectors for Ctrl treatment or combined Klf6+ Nr5a2 treatment. One week later, animals were challenged with pyramidotomy injuries, as described above. One-week post-injury, animals were euthanized, and the motor cortices were dissected. Dissected cortices were frozen on dry ice and stored at -80 degrees until library preparation. Frozen tissue was transferred to a pre-chilled 3 ml Kimble dounce homogenizer filled with 4ml Nuclei Lysis Buffer (Sigma, NUC101), supplemented with RNAase inhibitors (10mg/ml) (RNAase OUT, Thermofisher 10777019). Tissue was homogenized (20 strokes with pestle A and 25 stroked with pestle B) and allowed to incubate on ice for 5 mins. Following incubation, nuclei was centrifuged for 5 mins at 4 degrees (800G, low brake and acceleration) and the pellet was resuspended in 4ml Nuclei lysis buffer. Homogenate was incubated for 5 mins on ice. Following incubation, nuclei was centrifuged as described above and resuspended in 0.5ml 1X PBS with 1% BSA. Dissociated nuclei were filtered using a 20um filter and flow-sorted on a BD FACS Melody using an 80um nozzle to a goal of approximately 4500-5000 events (gatey strategy described above). Nuclei were sorted directly into 10X RT buffer (enzyme added only prior to GEM generation) and library preparation was performed on the 10X chromium platform according to manufacturer’s instructions (10x Genomics-Next GEM, Cat # PN1000121). Samples were sequenced at the University of Wisconsin-Madison Genomics core to a depth of ∼ 70,000 reads/nuclei (3000-4000 nuclei per library) on an Illumina NovaSeq platform. Raw FASTQ files were fed into the Cellranger pipeline (default parameter) described here - https://github.com/10XGenomics/cellranger. Datasets were then integrated using SEURATv3^39^and differential expression testing was performed according to default conditions (non-parametric Wilcoxon rank sum test).

### Code availability

There was no custom code development and all software used in data analyses are previously published, open-access and have been cited under the relevant methods section. Links to relevant software repositories/documentation is listed here -

WGCNA - https://horvath.genetics.ucla.edu/html/CoexpressionNetwork/Rpackages/WGCNA/; RNA-Seq analyses - https://github.com/ENCODE-DCC/rna-seq-pipeline;

EdgeR-https://www.bioconductor.org/packages/release/bioc/html/edgeR.html;

Activity-by-contact (ABC) -https://github.com/broadinstitute/ABC-Enhancer-Gene-Prediction; oPOSSUM 3.0-http://opossum.cisreg.ca/oPOSSUM3/;

Cytoscape - ClueGO - http://www.ici.upmc.fr/cluego/cluegoDocumentation.shtml; Harmonizome - https://amp.pharm.mssm.edu/Harmonizome/;

iRegulon - http://iregulon.aertslab.org/

Cellranger - https://github.com/10XGenomics/cellranger

SEURAT - https://github.com/satijalab/seurat

## Figure legends

**Supplemental Fig. 1** | **Expanded morphological screening data** (a) shows a transfected cortical neuron and tracing by cellomics, followed by schematic depictions of morphological parameters. (b-f) show the effect of forced expression of KLF6 alone, Klf6 with VP16-Stat3, or Klf6 with candidate TFs. In all cases measurements are normalized to those of within-plate control plasmids. Green bars indicate significant increases, and red bars indicate significant reductions compared to the values of klf6 alone (p<.05, 1-way ANOVA with post-hoc Fishers). A minimum of 150 individual cells from a minimum of three replicate experiments were used for each bar; full details are available in Supplementary Table 2.

**Supplemental Fig. 2** | **Co-injection of AAV2-Retro drives effective co-expression of transgenes in corticospinal tract neurons**. Adult mice received cortical injection of mixed AAV2-Retro-H2B-EGFP and AAV-H2B-mScarlet, followed two weeks later by injection of CTB-647 to cervical spinal cord. Animals were perfused three days later, and transverse sections of cortex examined by confocal microscopy. (a-d) show the distribution of CST neurons and AAV-expressed fluorophores. (e-h) show higher magnification views, illustrating efficient co-expression of both fluorophores in CST neurons (arrows). The circle indicates a rare instance of a CST neuron expressing EGFP (green, F) but not mScarlet (red, G). (I) quantifies the percent of transduced CST neurons that show single expression of either fluorophore or dual expression of both: 95.4% (± 1.51 SEM) of CST neurons were dually transfected. n=200 CST neurons scored from each of 5 mice. Scale bars are 1 mm (a-d) and 0.1mm (e-h).

**Supplementary Fig. 3** | **Summary of *in vivo CST* cross-midline growth after single or combinatorial expression of candidate TFs** (a) illustrates the unilateral pyramidotomy injury and indicates the approximate location of images. (b) shows a transverse section of spinal cord, eight weeks after pyramidotomy, with CST axons labeled by tdTomato. Vertical lines show the sampling regions in which axonal profiles were counted at 200, 400, and 600um from the midline. (c,d) provide example images from all animals in the two *in vivo* experiments; where images are missing, the reason for exclusion is indicated.

**Supplementary Fig. 4** | **Summary of tdTomato label in CST axons in the medulla, used to normalize CST counts in the spinal cord** (a) illustrates the unilateral pyramidotomy injury and indicates the approximate location of images. (b) shows a transverse section of an example medullary pyramid, nine weeks after cortical injection with AAV-TF treatment and AAV-tdTomato tracer, in which CST axons appear as red (tdTomato+) puncta as they intersect the plane of the section. The pyramid is outlined, and vertical boxes indicate the regions in which each individual axon was counted. Total axon numbers were estimated by multiplying axon counts by the total sampling area, divided by total medullary area. (c,d) provide example images from all animals in two *in vivo* experiments; where images are missing, the reason for exclusion is indicated.

**Supplementary Fig. 5** | **Summary of PKCγ signal in cervical spinal cord, used to verify unilateral ablation of the CST** (a) illustrates the unilateral pyramidotomy injury and indicates the approximate location of images. (b) shows a transverse section of cervical spinal cord, eight weeks after pyramidotomy injury, with PKCγ signal (green) readily detectable in the intact but not transected CST (white arrow). (c,d) provide example images of the dorsal columns after PKCγ staining from all animals in two *in vivo* experiments; where images are missing, the reason for exclusion is indicated.

**Supplementary Fig. 6** | **RNAscope and immunohistochemistry confirm expression of candidate transcription factors** (a-e) show coronal section of adult mouse cortex, eight weeks after cortical injection of AAV expressing candidate transcription factors, with arrows marking the site of injection. RNAscope or appropriate antibodies were used to probe for expression. RNAscope signal from probes directed against the expressed TFs is present at the injection site, which at higher magnification (a’-d’) shows the characteristic punctate detection of transcripts. e and e’ show detection of Eomes by immunohistochemistry. (f-u) Adult mice received cortical injection of AAV-Klf6, AAV-Nr5a2, and AAV-tdTomato at 1.5:1.5:1 ratio, the same used in axon growth experiments. Two weeks later cortices were examined by fluorescent in situ hybridization (RNAscope) to visualize expression of Klf6 and Nr5a2. (f-h) show tdTomato at the site of viral injection, (i-k) show Klf6 expression, (l-n) show Nr5a2, and (o-q) show the overlay. Note that tissue distant from the injection site displays very low levels of Klf6 and Nr5a2 detection, while virally-expressed transgenes are readily detected at the site of injection. (r-t) show a cortex that received AAV-tdTomato and AAV-Cre control (arrowhead), with low detection of endogenous Klf6 (s) and Nr5a2 (t) transcripts. (u) tdTomato+ cells were classified according to dual, single, or no expression of Klf6 and Nr5a2 transcripts; more than 90% of tdTomato+ cells expressed both transcripts. n = 475 cells analyzed from three animals. Scale bars are 1mm (a-e, f, i, l, o, r-t), 50 µm (a’-e’), and 100 µm (g, h,j, k, m, n, p, q).

**Supplemental Fig. 7** | **Cortical injection of AAV-Klf6 and AAV-Nr5a2 does not elevate gliosis, inflammation, or cell death compared to injection of control AAV** (a-d) Adult mice received cortical injection of AAV-tdTomato with AAV-Cre or combined AAV-Klf6/AAV-Nr5a2 (KN). Three days later, animals were sacrificed and coronal sections of cortex examined for gliosis (GFAP), inflammation (Cdllb), or cell death (TUNEL). Both control and KN-injected animals showed similar levels of GFAP and cd11b near the site of injection (a, b) and minimal cell death (c,d). (e-h) Animals received cortical injection of AAV-Cre control or AAV-K/N and cervical injection of CTB-647 to label CST neurons. Four weeks after viral injection coronal sections of cortex were stained for GFAP, CD11B, or TUNEL reactivity. In both control and KN-treated animals CST neurons were apparant in the vicinity of the injection, and minimal levels of GFAP and CD11B persisted. (g,h) TUNEL signal was rare in both treatments, and never co-localized with CST neurons. (i) Shows adult cortex 2 days after injection of Staurosporine. Numerous cells are TUNEL-positive (green), confirming assay sensitivity. (j) quantifies the average number of TUNEL+ cells located within 500µm of the injection track. Compared to staurosporine, AAV-Cre control and AAV-KLF6/Nr5a2 produce TUNEL+ cells at low numbers that do not statistically differ (p>.03, 1-way ANOVA with post-hoc Dunnett’s). n=3 animals in each group, three replicate sections per animal. Scale bars are 1mm (a-h) or 100 µm (i).

**Supplementary Fig. 8** | **Single nuclei RNASeq analyses validate transcriptional changes following combined KLF6/Nr5a2 gene treatment** (a) Overview of sample collection for Single nuclei RNA-Seq analysis. (b) UMAP visualization of 3383 nuclei (Control) and 3038 nuclei (KLF6+Nr5a2 treated) that passed QC filtering (see methods) confirms qualitative concordance across groups (c) Expression of key marker genes delineated nuclei clusters specific to Corticospinal tract neurons (d) Bioinformatic analyses confirmed ∼ 50% overlap in Klf6/Nr5a2 responsive target genes between the two independent RNA-Seq approaches. Regulatory network analysis of genes upregulated after combined Klf6/Nr5a2 overexpression confirmed sub-networks enriched for functional categories relevant to axon growth (e-g) Agreement of log2 Fold change in expression of functionally distinct Klf6/Nr5a2 responsive target genes between the two independent RNA-Seq approaches. Differential testing – Non-parametric Wilcoxon rank sum test (SEURAT v3). n=3 animals/rep and 2 reps/treatment.

## Acknowledgments

This work was supported by grants from NINDS (5R01NS083983, R21NS106309), The Craig Neilsen Foundation, The Bryon Riesch Paralysis Foundation, and computational allocations from NSF-XSEDE. We wish to thank Erik Van Newenhizen for technical assistance. The authors acknowledge ENCODE consortia for generating the developmental time-series NGS datasets used in this study. The authors acknowledge the use of Biorender to generate illustrations and graphics used in the manuscript. The authors declare no competing financial interests.

